# Vision-to-value transformations in artificial neural networks and human brain

**DOI:** 10.1101/2021.03.18.435929

**Authors:** Trung Quang Pham, Takaaki Yoshimoto, Haruki Niwa, Haruka K Takahashi, Ryutaro Uchiyama, Teppei Matsui, Adam K Anderson, Norihiro Sadato, Junichi Chikazoe

## Abstract

Humans and now computers can derive subjective valuations from sensory events although such transformation process is essentially unknown. In this study, we elucidated unknown neural mechanisms by comparing convolutional neural networks (CNNs) to their corresponding representations in humans. Specifically, we optimized CNNs to predict aesthetic valuations of paintings and examined the relationship between the CNN representations and brain activity via multivoxel pattern analysis. Primary visual cortex and higher association cortex activities were similar to computations in shallow CNN layers and deeper layers, respectively. The vision-to-value transformation is hence proved to be a hierarchical process which is consistent with the principal gradient that connects unimodal to transmodal brain regions (i.e. default mode network). The activity of the frontal and parietal cortices was approximated by goal-driven CNN. Consequently, representations of the hidden layers of CNNs can be understood and visualized by their correspondence with brain activity–facilitating parallels between artificial intelligence and neuroscience.

## Introduction

Valuation is an important mental process for decision-making. Subjective valuation is supported by sensory stimuli such as somatosensory, auditive, and especially visual input and is performed consciously or subconsciously. A recent study [1] has revealed that visual/gustatory stimuli can be simultaneously converted into categorical and value-related information. Value-related information has been mainly related to activity on the orbitofrontal cortex and found to be independent of sensory modalities. Several anatomical studies have suggested an occipitotemporal–orbitofrontal pathway connecting the inferior temporal cortex, as the end of the ventral visual pathway, with the orbitofrontal cortex [2, 3, 4]. However, the neural mechanism to transform visual stimuli into value-related information is largely a blackbox.

The categorical transformation of visual information (i.e., vision-to-category transformation) has been extensively investigated due to the emergence of artificial neural networks (ANNs), which provide unique architectures consisting of nodes stacked into consecutive blocks to resemble the sequential computations that occur in individual neurons and regions of the human brain. Another resemblance of an ANN to the human brain is the use of nonlinear activation as well as operations of pooling, normalization, convolution, among others to perform computations [5]. Theoretically, an ANN can provide a mechanism for projecting any suitable input to a desired output if provided with sufficient data and appropriate instructions [6]. A feedforward ANN inherently models a sequential computation. Thus, goal-driven approaches have been devised to first optimize the ANN parameters to solve behavioral tasks [5, 7], and then use the ANN as a simplified brain model for further investigation. For instance, the convolutional neural network (CNN), a predominant ANN for image classification in computer vision, has succesfully predicted the neural responses in the visual V4 and inferior temporal (IT) cortex [8], and captured the hierarchical organization of voxel responses in functional Magnetic Response Imaging (fMRI) across the visual ventral stream [9].

Functional processing hierarchies have been found in various cognitive systems of the brain [10, 11, 12]. Margulies et al. [13] revealed a global gradient that connects the primary sensorimotor regions and the transmodal regions in the default mode network (DMN). The global gradient is called the principal gradient (PG) because it accounts for the highest variability in human resting-state functional connectivity. A meta-analysis using the NeuroSynth database [14] further suggested the relationship between the PG score and cognitive functions. A low PG score is associated with sensory perception and motion (unimodal regions) whereas a higher score is associated with the high-order abstract and memory-related processes, such as emotion and social recognition (transmodal regions). Therefore, the transformation of visual information into subjective valuations (i.e., vision-to-value transformation) may be a hierarchical process along the PG.

While previous studies typically assumed the transformation to be consistent across individuals, as in the case of object classification, that is not the case for vision-to-value transformation because of the high variance in aesthetic preference. Hence, there remains a need for individually optimized CNNs. In this study, we developed a group of CNNs that are capable of predicting the subjective valuations of art pieces (oil paintings) by various participants (one CNN per individual). Our motivation is to use the intrinsic hierarchy of the CNN (i.e., the output of each layer is the input of the next one) to unveil a hierarchical structure during vision-to-value transformation in the human brain. We then constructed maps between the brain activity and the CNN computations by performing the representational similarity analysis between fMRI signals and the output of the CNN hidden layers. Based on these maps, we assigned a PG score to every voxel contained within a region that corresponds to each CNN layer. Then we investigated the correlation between the arrangement of the CNN layers and the PG. By specifying correspondence between representations of the respective hidden layer and brain regions, we obtained visual representations of the hidden layer computations.

## Results

### CNN optimization per participant

To optimize the CNN to reflect the aesthetic preferences of each participant, we applied transfer learning as illustrated in Fig. 1a. First, a baseline CNN was built using the VGG-16 architecture, a contest-winning CNN, with batch normalization (see section Materials and Methods in the Supplementary Materials for details). As the original VGG-16 structure is pre-trained on the ImageNet dataset for object classification, the baseline CNN was modified by transfer learning to perform regression on an art auction dataset (38,059 art items with hammer prices) that was retrieved from the LiveAuctioneers (https://www.liveauctioneers.com/). The baseline CNN achieved a median Spearman correlation coefficient of 0.41 and 0.34 on the validation (n = 3054) and test (n = 7517) sets, respectively. The Spearman correlation was used because it is less sensitive to the outliners in the dataset.

**Figure 1:**
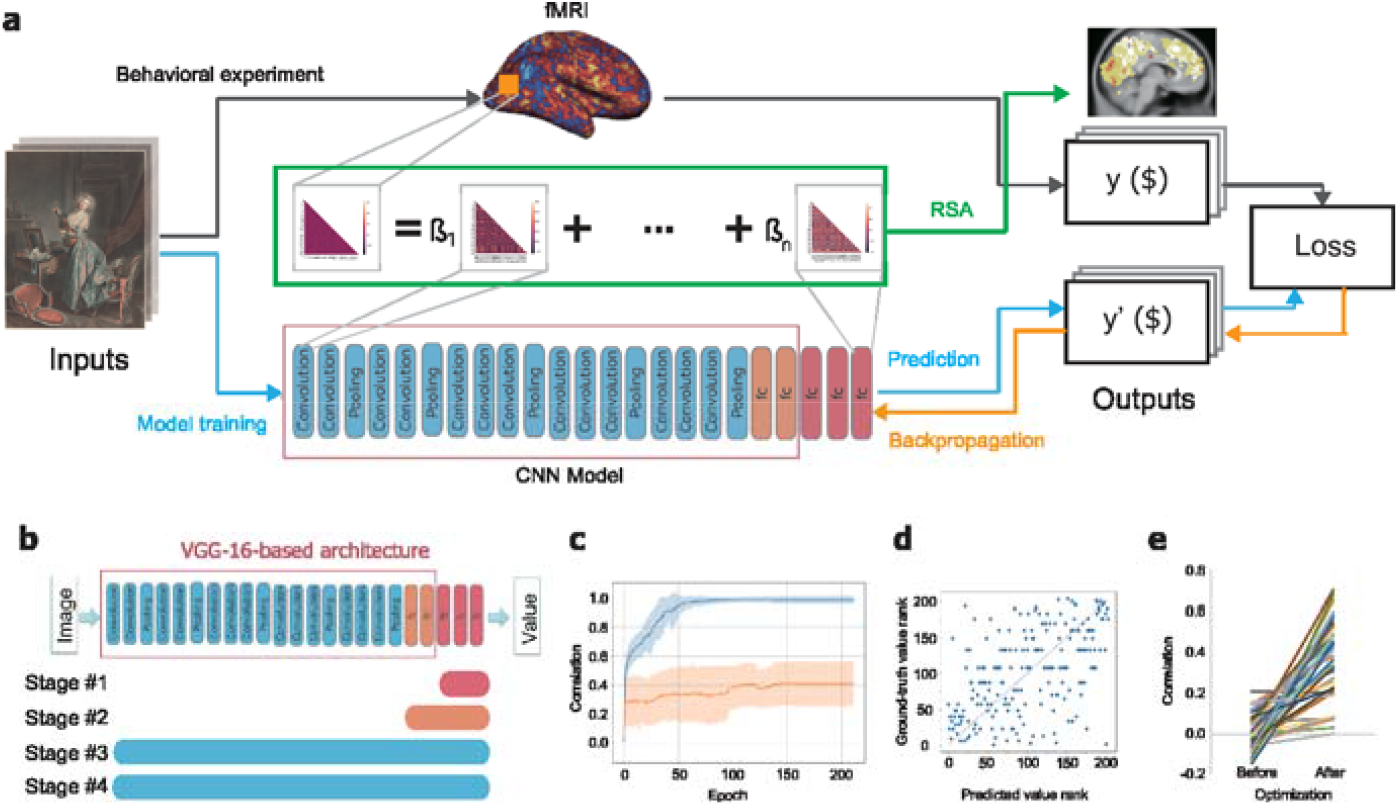
Optimization of the CNN to reflect the aesthetic preferences of each participant. (**a**) Overview of the CNN development and analysis. Black arrows indicate a behavioral data during fMRI to obtain ground truth values (y). Blue arrows indicate the computation of the CNN model on each input to obtain predicted values (y’). Orange arrows indicate backpropagation for updating the parameters of the CNN model. The green arrow indicates representational similarity analysis (RSA). β indicates the coefficient for each CNN representational similarity matrix. (**b**) Four stages of transfer learning in sequential order. At each stage, a portion of the model (bar) was trained. (**c**) Change in correlation between ground truth and predicted values as a function of training epochs of a typical, on training (blue) and validation (red) datasets. (**d**) Plot of the rank of ground truth values as a function of the rank of predicted values of a typical IO-CNN, on the validation dataset. (**e**) Improvement of all IO-CNN Spearman correlations on the test dataset, before and after individual optimization.

Then, for each of the 37 participants, we obtained the corresponding individually optimized CNN (IO-CNN), which was derived from the baseline CNN but subject to additional transfer learning over the valuation data of each participant. The valuation data were obtained during an fMRI experiment in which the participants were asked to quote their prices for the art pieces. The additional training based on transfer learning was conducted over four stages (Fig. 1b). During the first three stages, the layers were gradually unfrozen, starting from the last three fully connected layers followed by all the fully connected layers and finally all the layers (see section Materials and Methods for details). Figure 1c shows the learning curve of a typical IO-CNN (110 epochs in total) and Fig. 1d shows the correlation between the predicted and ground truth values (Spearman correlation coefficient *r*=0.53 on the test set). The IO-CNNs suitably predicted the participant’s individual aesthetic preferences, with a median correlation coefficient of 0.35 on the test set. From 37 networks, 33 had a correlation coefficient higher than 0.15 (*p*<0.05) (Fig. 1e). Obviously, the baseline CNN without individual optimization showed a poor predictive power (median Spearman correlation coefficient of 0.0016) across all subjects on the test set, justifying the individual CNN optimization to predict aesthetic preferences.

### Mapping representational correspondences between IO-CNNs and brain regions

To examine the representational correspondence between each layer of the IO-CNN and brain regions, we conducted a representational similarity analysis [15]. Two kinds of correlation matrices were constructed, one for neural activity at every searchlight (33 voxels) and one for every layer of the IO-CNN (Fig. 1a). The correlation matrix of neural activity across the art images at a given searchlight was transformed into a vector. Then, the vector was subjected to a multivariate linear regression considering the vectors from the correlation matrices of the IO-CNN layer across images as independent variables. The regression coefficients were computed for each layer and then subjected to one sample t-tests across participants. We found that early visual areas (e.g., V1 and V2) are associated with shallow IO-CNN layers (Fig. 2), whereas higher areas in the visual cortex (e.g., the parahippocampal and fusiform gyri) are associated with deeper IO-CNN layers (Fig. 2a). Throughout the paper, all the correspondence maps were overlaid by Gordon’s parcellation [16] for better visualizations. These findings are consistent with previous studies that employed CNNs for object recognition [9, 17, 18]. Furthermore, the valuation information (i.e. in layer FC 18) obtained in this study is related to activity in multiple brain regions such as the ventromedial prefrontal cortex and lateral orbitofrontal cortex, also being consistent with the previous studies [1, 19]. These value representations are not limited to â€œhedonic hotspotsâ€ such as the orbitofrontal cortex and insula [20], but they also covered broad regions of the DMN [21]. Importantly, we found that broad regions in the prefrontal and parietal cortices are associated with the intermediate representations between visual and valuation information, rather than with valuation exclusively. To visualize these layer-to-region relationships, we numbered the layers consecutively from 1 (Conv 1) to 18 (FC 18), assigned a colormap to the number sequence and highlighted the voxel with the highest association to the layer activation, obtaining the results shown in Fig. 2b.

**Figure 2:**
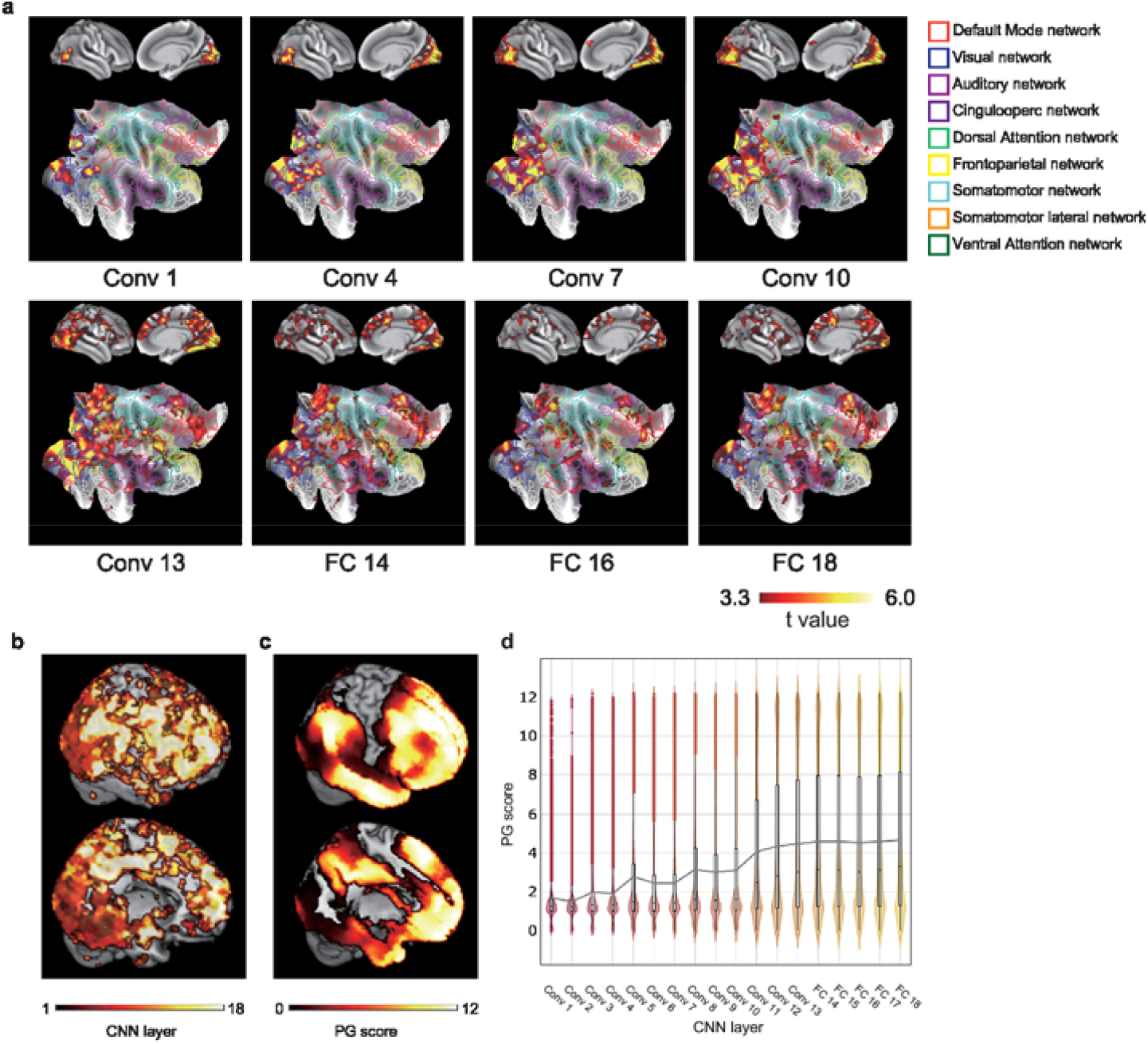
Brain–CNN correspondence maps. (**a**) Correspondence maps between the neural network’s layer activation and the brain’s voxel activation (Conv, convolution layer; FC, fully connected layer). The background image was overlaid by 9 networks from [16] for better visualizations. (**b**) Summary map of brain-CNN correspondence (upper: lateral view, lower: medial view). Each voxel was highlighted with the number sequence of it’s highest associated layer. (**c**) The PG score arranges the cortex into a continuous spectrum from the primary sensory regions to the DMN. (**d**) The correlation between the PG score and CNN sequential layers. The violin plots represents the distributions of PG score. The boxes span the first to third quartiles. The black dot inside the boxes represents the median. The gray line connects all the means of the boxes.

The correspondence maps show that the activity corresponding to deeper CNN layers gradually shifted from visual areas to the DMN. This pattern resembles the PG of cortical connectivity [13], which is the primary axis of an intrinsic coordinate system characterizing the cortical architecture and bound by primary sensorimotor areas and transmodal areas. To analyze the representational IO-CNN gradients according to the PG, we assigned color-coded PG scores to all voxels contained within a region corresponding to each IO-CNN layer. A high score indicates that the voxel distribution over the cortex is closer to the transmodal PG pole (i.e., the DMN), and a low score indicates a distribution close to the sensory pole (Fig. 2c). The PG scores for all voxels were obtained from Margulies et al. [13](https://www.neuroconnlab.org/data/index.html). Figure 2b–d show the sequential arrangement of the IO-CNN layers is highly correlated with the PG score (*r*=0.25; *p*<0.001) (Fig. 2d). The significant positive correlation between the PG score and the CNN sequential layers indicates that the pathway for the vision-to-value transformation revealed by the IO-CNNs is aligned with the PG. Therefore, concrete, unimodal sensory information is likely to be sequentially transformed into abstract, transmodal cognitive representations through hierarchical processing along the PG. Although such a hierarchical structure was implied by the meta-analysis in [13] and studies on intracranial electric stimulation [22] and task-related fMRI scans based on encoding models [23, 24], we are the first to provide evidence for the sequential processing along the PG. These results also indicate that the hierarchical vision-to-value transformation is a global process involving areas on the whole brain.

To further evaluate the existence of a hierarchical processing structure on the cerebral cortex, we counted the number of significant voxels per CNN layer in various regions of interest (Fig. 3). As expected, the primary visual cortex (V1) is associated with shallow layers, such as Conv 1 and 2. Interestingly, multiple regions, including anterior inferior frontal sulcus (Fig. 3d), area posterior 32 (p32) (Fig. 3e), and 31a (Fig. 3f) are associated with valuation, while their surrounding regions, including area anterior 9-46v (a9-46v) (Fig. 3d), 10 dorsal (10d) (Fig. 3e), and 23d (Fig. 3f) are associated with the intermediate representations between vision and valuation. These results further confirm that vision is transformed into valuation via hierarchical processing on the parietal and frontal cortices.

**Figure 3:**
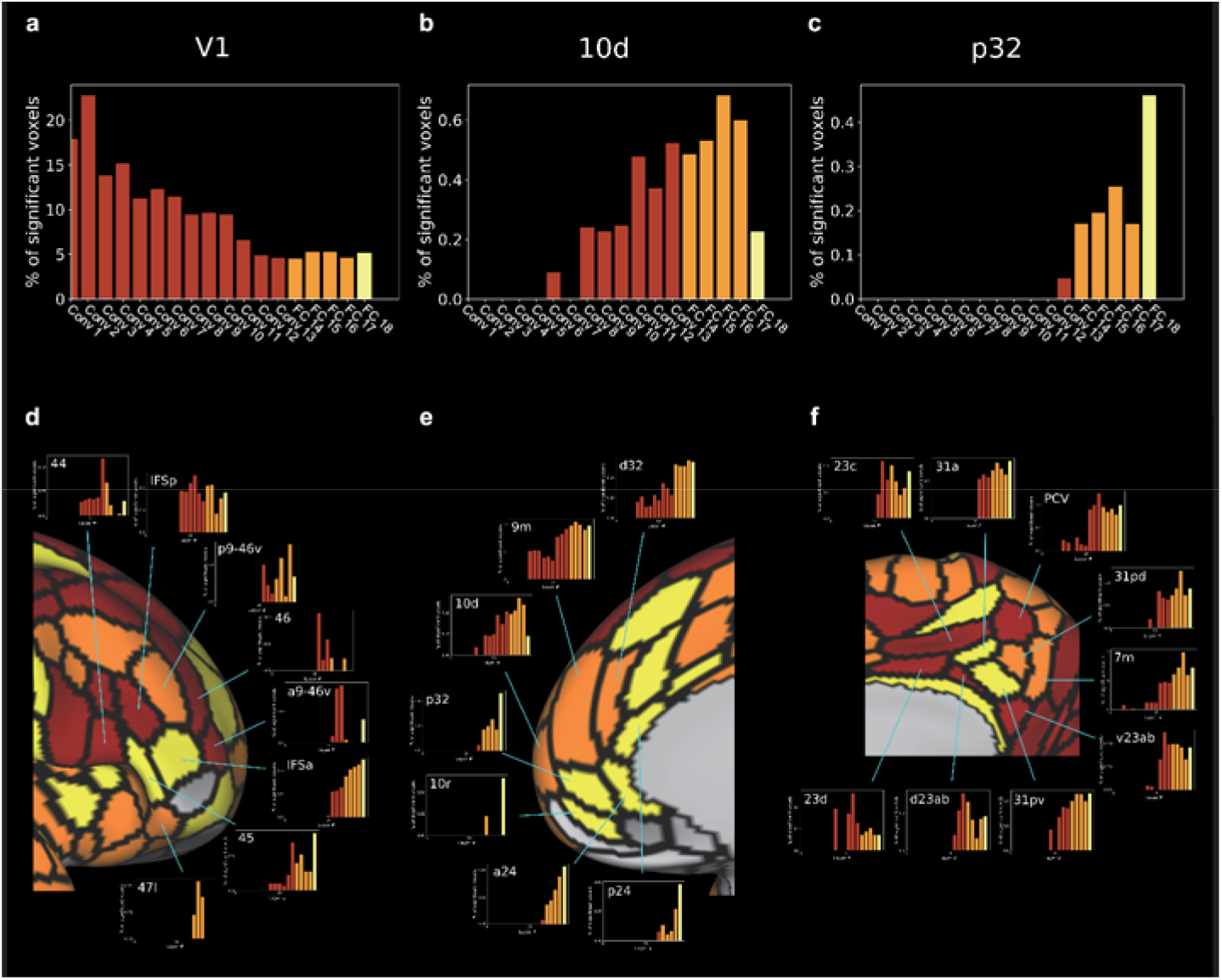
Brain–CNN correspondence within ROIs. (**a–c**) The proportion of significant voxels in the group analysis of brain–CNN correspondence for each layer in V1 (a), 10d (b), p32 (c). The color of a bar indicates the depth of the layer (yellow: FC 18, orange: FC 14–17, red: Conv 1–13). (**d–f**) Visualization of the difference in brain–CNN correspondence across regions in the lateral prefrontal cortex (d), medial prefrontal cortex (e) and medial parietal cortex (f). The layer color showing the highest proportion was given to each of the HCP MMP ROIs. For detail list of abbreviations, see Table **Error! Reference source not found**..

A high variability regarding appreciation of beauty (i.e., valuation) may be expected between participants. For measuring the reliability of valuation, we analyzed the reproducibility of ratings for the same paintings. The participants completed a reproducibility check in a behavioral experiment, where they were asked to rate the same art piece twice in an interval of approximately 2 hours. We then calculated the correlation between these two scores per participant (median *r*=0.68). Based on this reliability score, we split the participants into reliable and unreliable groups (Fig. **Error! Reference source not found**.). The participant showing the median correlation was excluded. Figure 4a and b show that the voxels in the frontal and parietal cortices correspond to deep IO-CNN layers in reliable participants but not in the unreliable ones. While the ventral visual pathway were automatically included regardless of participant’s reliability scores, the further pathway of transformation were vigorously activated in reliable ones.

**Figure 4:**
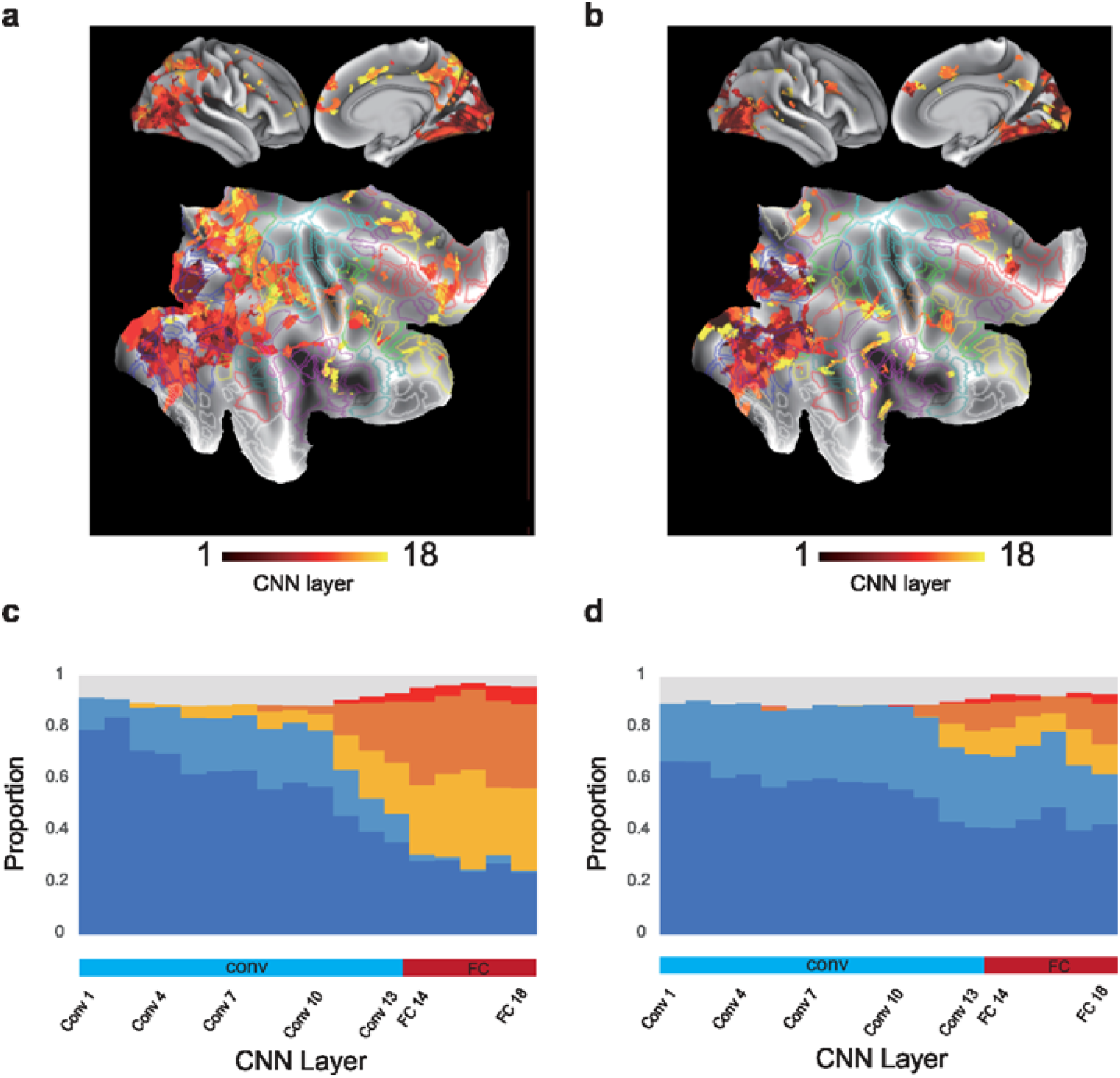
Comparison of Brain–CNN correspondence maps between reliable and unreliable participants. (**a–b**) The brain–CNN correspondence map for reliable group (a) and unreliable group (b). N=18. (**c–d**) The layer-wise proportion of significant voxels in the occipital (dark blue), temporal (blue), parietal (gold), frontal (orange), insula and cingulate (red) and other cortices for the reliable group (c) and unreliable group (d).

To elucidate the difference in brain–CNN correspondence between the reliable and unreliable groups, we obtained the layer-wise proportion of brain–CNN correspondence on the occipital, temporal, parietal and frontal cortices as well as that on the insula and cingulate cortex for the reliable (Fig. 4c) and unreliable (Fig. 4d) groups. The transition from visual to transmodal areas is more prominent in participants from the reliable group. In these participants, the ventral visual pathway consisting of the occipital and temporal cortices was first recruited, followed by the frontal and parietal cortices were recruited for further computation (Fig. 4c). In contrast, for the unreliable group, even the processing close to the valuation output (i.e. FC 14–18) relies primarily on the ventral visual pathway but not on the frontal and parietal cortices (Fig. 4d). The IO-CNNs in both the reliable and unreliable groups performed comparably well on their training dataset, which contains the art-prices pairs obtained from the fMRI experiment. Hence, the prominent difference between these groups on the frontal and parietal cortices indicates the crucial role of these brain areas for reliable vision-to-value transformation from object to value.

### Brain–CNN correspondence for interpreting CNN representations

A major drawback of deep ANNs (including CNNs) is the difficulty to understand their inner computations. We demonstrate the use of brain–CNN correspondence to visualize internal CNN representations and improve interpretability. Figure **Error! Reference source not found**. shows the correlation between the actual and predicted valuations according to the individual CNN optimization. Although a suitable prediction on the test dataset was achieved at the first training stage, the further optimization slightly improved the prediction in Fig. **Error! Reference source not found**.b. At early stages of CNN training, only voxels in the occipital and temporal cortices correspond to CNN representations (Fig. 5a and 5b). As training proceeds, the CNN representations, especially those of the fully connected layers, become similar to the representations of the frontal and parietal cortices. Notably, the representations are substantially altered during training stage 3 despite the negligible prediction improvement of aesthetic preferences on the test dataset. Thus, the CNN prediction performance does not fully reflect the change in the representation in the hidden CNN layers.

**Figure 5:**
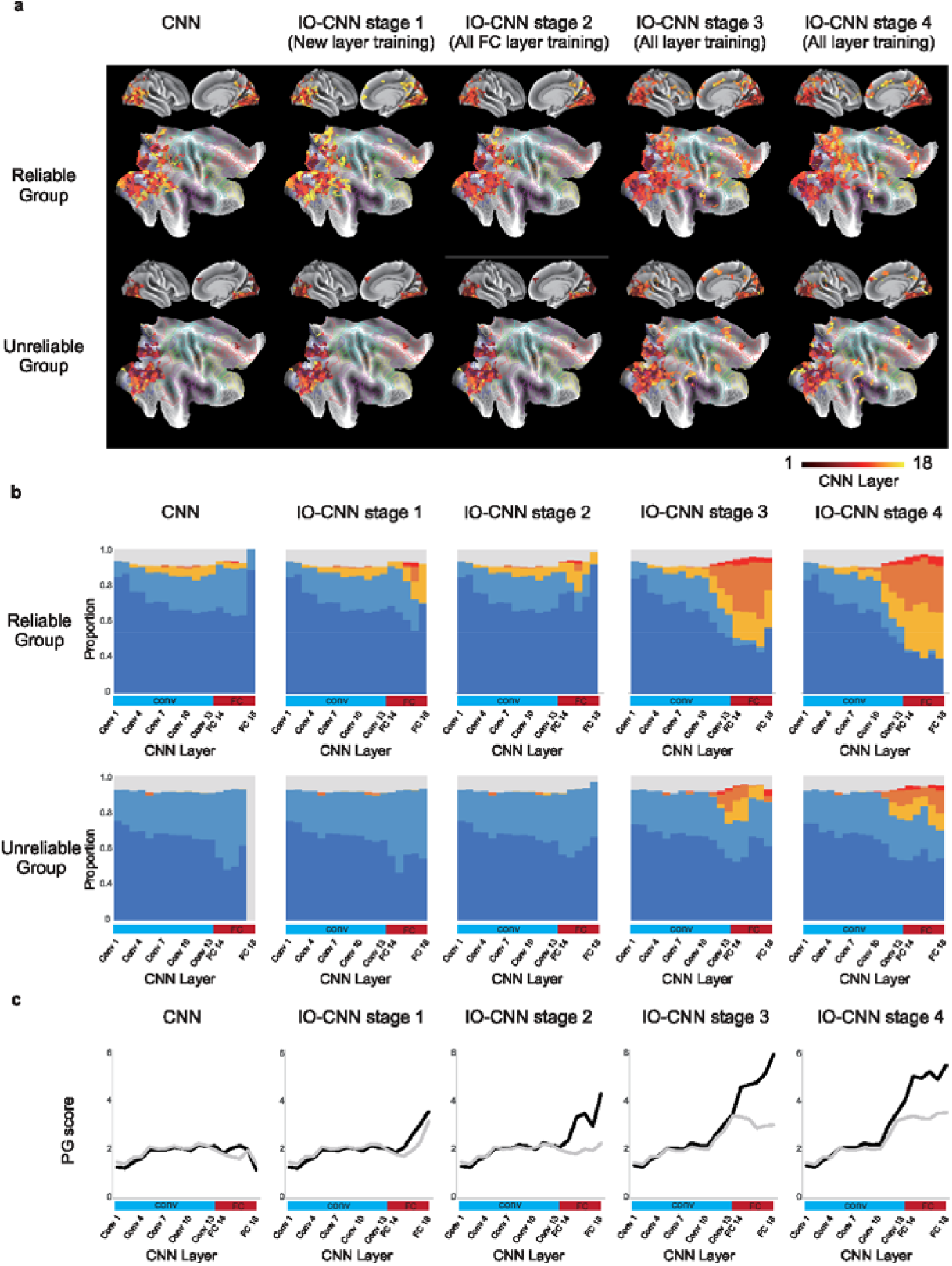
Visualizing change in CNN representations over individual optimization. (**a**) Brain–CNN correspondence maps underwent reconfiguration after each optimization stage. The colors are encoded in similar manner as in Fig. 2a. (**b**) The layer-wise proportion of significant voxels for the reliable and unreliable groups. The colors are encoded in similar manner as in Fig. 4c–d. (**c**) Mean PG score for brain regions corresponding to each layer at each training stage (gray: unreliable group, black: reliable group). The reliable group showed more pronounced variation in PG scores across corresponding layers, particularly in fully connected layers (FC 14–18).

To statistically evaluate the change in CNN representations through individual optimization, we calculated the PG scores for the reliable and unreliable groups at every optimization stage (Fig. 5c). The CNN representations became increasingly more aligned with the hierarchical structure across the human brain along with the improving CNN prediction accuracy for individual aesthetic preferences (Table **Error! Reference source not found**.). Furthermore, in the brain regions corresponding to fully connected layers (FC 14–18), the PG scores gradually increased as training proceeded. Given that these PG scores are strongly associated with a spectrum of concrete-to-abstract cognitive domains [13], our results indicate that CNN representations in higher layers become increasingly abstract and remote from sensory information as individual optimization proceeds.

Finally, to confirm that the layer–PG correspondence is shared in the internal representations across IO-CNNs, we obtained the correspondence after replacing the baseline VGG-16 by three popular CNN architectures, namely, DenseNet [25], ResNet [26], and Inception Network [27] (Fig. **Error! Reference source not found**.). These architectures also achieve high layer–PG correspondence (Fig. 6; Table **Error! Reference source not found**.),indicating that hierarchical sequential computations similar to the PG may be characteristic across families of CNNs. Overall, these results suggest that examining relations between CNN computations and brain activity is useful for investigating the internal CNN representations regardless of the specific CNN architecture.

**Figure 6:**
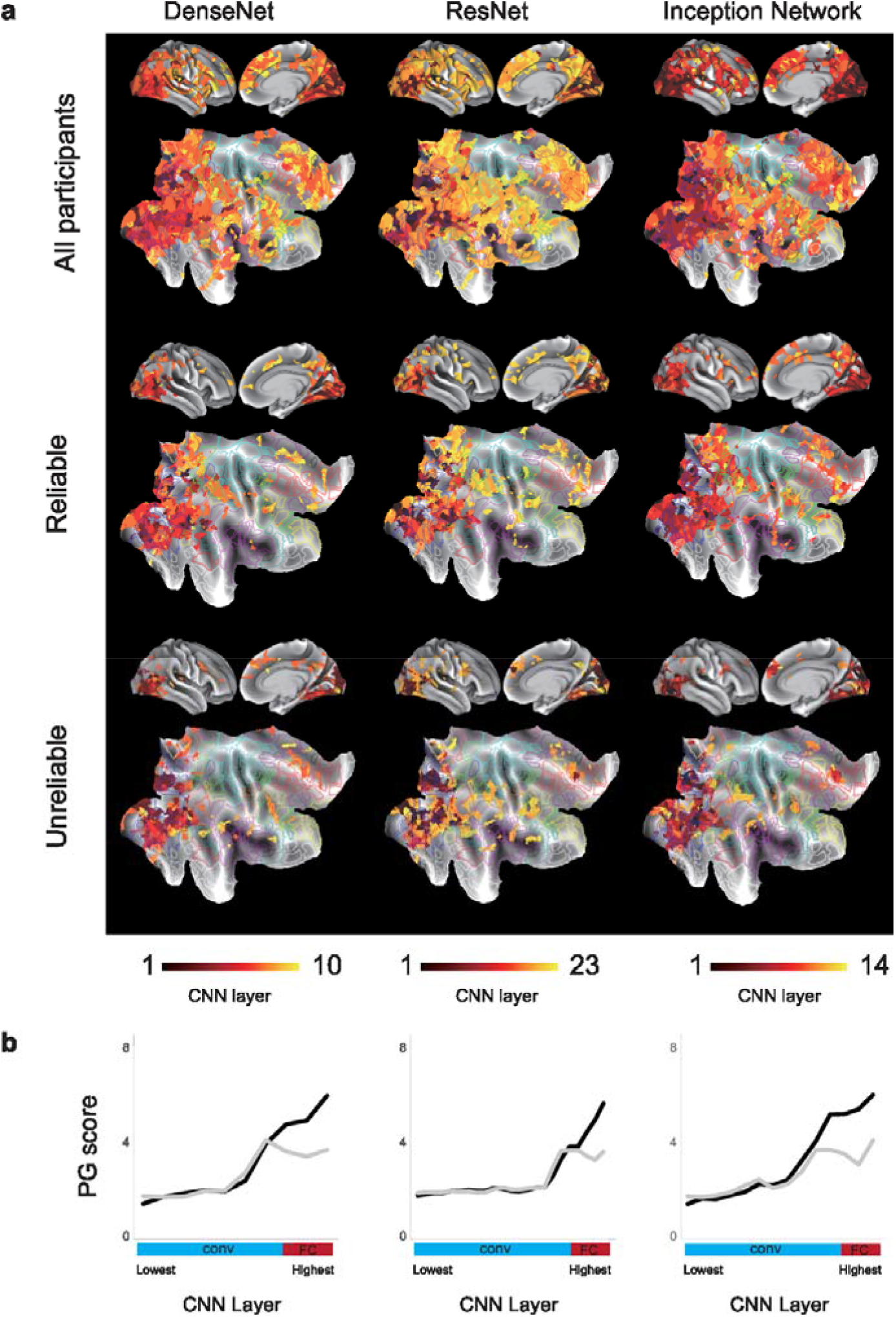
Differences in brain–CNN correspondence across 3 different CNN models. (**a**) Brain–CNN maps of DenseNet, ResNet and Inception Network, from left to right respectively. The colors are encoded in similar manner as in Fig. 2a. (**b**) Mean PG scores for reliable (black) and unreliable (gray) groups, across three CNN architectures.

## Discussion

In this study, we demonstrated the usefulness of IO-CNNs to investigate the hierarchical processing during vision-to-value transformation. Although the correlation between the IO-CNN predicted values and ground-truth values varied between the participants, it was generally improved compared with the baseline CNN (Fig. 1e). By analyzing the correspondence between the IO-CNN and human brain activity, we unveiled a hierarchy of vision-to-value transformation in the brain as a gradient-guided process from the visual area to the DMN (Fig. 2). The brain–CNN correspondence maps were not biased toward a hierarchical structure by using ANNs because the assignment of voxels to the CNN layers was conducted independently of the network architecture via multivariate linear regression. The hierarchical vision-to-value transformation revealed by the IO-CNNs is aligned with the PG (Fig. 3). We also found a high CNN layer–PG correspondence for reliable participants compared with unreliable participants. We observed the most prominent difference in the frontal and parietal cortices, indicating that reliable vision-to-value transformation is supported by the activity in transmodal regions. These results were reproduced across various CNN architectures including VGG-16, DenseNet, ResNet, and Inception Network, suggesting that hierarchical sequential computations similar to the PG are a shared characteristic between CNNs. Overall, our results indicate that valuation is the result of the gradual transformation of external information through the PG pathway, which begins with sensory representations and ends in DMN processes.

### Value representations over the sensation-to-value gradient

In a previous work [1], we showed that information from multiple sensory modalities is converted into a common supra-modal valuation code on the medial and lateral orbitofrontal cortex. This organization mode suggests that gradients of representational structures exist along sensation-to-value pathways for each sensory modality, where low- or mid-level sensory features are gradually integrated with valuation-relevant information throughout processing. Given the current knowledge about the brain structure and function, this sort of gradual representation integration is more reasonable than a sharp modular division of labor between sensory areas and valuation-related areas of the brain [28]. Using IO-CNNs as models, we reached findings consistent with such processing pathway in the present study.

In future research, the levels of the processing hierarchy that elicit neural representations of valuation should be identified. In addition, the variations of these levels according to the task demands and learning should be determined. Although our results suggest an abstract representational format distributed over the DMN, these representations are not always required for tasks involving valuation. We may explore heuristic strategies to obtain efficient solutions to a value-inference problem through matching with familiar sensory representations. Such strategies are expected to be consistent with the proposal of Kahneman [29] in his cognitive heuristic of substitution. For instance, if a person is asked how much money they would like to contribute to save an endangered species, they are likely to decide based on an emotional response to ideas such as a dying dolphin rather than on a cost–benefit calculation. In the absence of a reliable or efficient way to estimate subjective valuation in abstract terms, a value-laden sensory image may serve as a concrete surrogate.

The notion that abstract mental representations are grounded in sensorimotor imagery has a long history in the cognitive sciences [30] and is supported by evidence from human neuroimaging studies [31], mainly related to concepts and language. We expect that an analogous organization may characterize valuation. As our findings suggest that the transformation from sensory input into abstract value unfolds gradually across the PG, the human brain is likely to exploit intermediate value representations that retain sensorimotor features. This processing strategy may play an important role in cultural responses such as the appreciation of visual arts and other behaviors. More generally, this strategy suggests that sensory experience is influenced by emotions before any subsequent appraisal [32].

### Role of DMN in functional organization of valuation

Although previous studies have mainly focused on the representation of valuation in the orbitofrontal cortex [1, 33, 34, 35], we found multiple representations of valuation across several areas associated with the DMN, such as the temporoparietal junction, parietal cortex, and ventromedial prefrontal cortex (Fig. 2). Thus, DMN activity may be essential during valuation. There may also be a more intrinsic relationship between valuation and the DMN.

Various studies have demonstrated an immediate encoding of value information in areas coinciding with the DMN, and coordinate-based meta-analyses of the brain networks involved in valuation have indicated significant overlap with the DMN, particularly in areas such as the ventromedial prefrontal cortex and posterior cingulate cortex [36, 37]. Acikalin et al. [36] suggested that there may be an underexplored link between the computational function of the â€œsimulation of internalized experienceâ€ and valuation. This finding complements recent developments in reinforcement learning that go beyond the classical emphasis on motor sequences by uncovering strategic interactions between the reinforcement learning system and cognitive operations such as episodic memory [38] and prospection [39], which are commonly associated with the DMN. Although the DMN has often been characterized as a â€œtask-negative networkâ€ that deploys neural resources toward internally oriented cognition at the expense of engagement with the external world, the DMN processes may be employed for the realization of external goals through coupling either with striatal reinforcement learning or with functional networks that guide action [40, 41]. In our behavioral experiment, the participants received novel visual stimuli from the environment and projected this information onto a subjective measurement space that registers a personal value judgment. The task thus merged externally and internally oriented processes, requiring the activation of the PG [13] across its full spectrum, from early sensory features to abstract transmodal representations that encode internalized experience. Hence, we speculate that the PG may represent an external–internal cognition axis.

Furthermore, several recent studies have found evidence of an intrinsic link between DMN activity and aesthetic appreciation [42, 43], suggesting that the DMN serves as a core for assessing domain-general aesthetic appeal. Our results support this finding and extend it by identifying the DMN as the functional locus. Across both anatomical and representational levels, our correspondence maps show the pathway over which visual input is transformed into value judgments, thus offering insights into the architecture and brain-wide dynamics of aesthetic experience. The richness of CNN models also allow to model individual differences in representational structures that underpin variations in aesthetic preferences.

### Variation in representational structure over time

The high performance of the IO-CNNs maintained for both the 200 (fMRI experiment) and 400 (behavioral experiment) art-piece datasets indicates that these models successfully captured the individual aesthetic preferences of each participant. The experiment trials were performed on different days, and although some participants were more reliable than others in reproducing their initial valuations, the overall preferences did not shift in any systematic way over time.

Over longer time frames, participants may shift their aesthetic preferences due to either new external information or internal changes. If we retrain the IO-CNNs over a longer period by performing transfer learning with new data, it would be possible to analyze the corresponding variations in intermediate representations. Given the potential of our method to capture rich individual differences among participants, determining individual trajectories of representational variation over time is a promising research area. The methods in this study can be employed to improve our understanding, both across individuals and over time, about the variability in representational structures that explains changes in valuation. By mapping these representations onto brain regions, our method may also provide insights about known mechanisms of neural plasticity mediating these changes.

### ANNs for investigating brain function

Our findings in this study demonstrate the feasibility to extract the novel insights about brain mechanisms by applying supervised learning to IO-CNNs tailored for different individuals. The IO-CNNs can describe the intrinsic transition of cognitive processing from perception of the external environment to the monitoring of internal states, converging with the functional organization of the cortex and its connectivity PG [13]. Hence, our study extends the domain of utility of CNNs compared with previous works [8, 9, 44].

There are various limitations that remain to be addressed regarding the use of CNNs to analyze brain functions. First, interpretability is limited due to the lack of a comprehensive method for both analyzing deep ANNs and understanding internal states of the brain. Changes in the similarity matrix occurred across all CNN layers per trial, and regions of interest in the brain could be visualized using the representational similarity analysis. However, the exact intermediate representations per trial remain difficult to describe. Second, training a group of IO-CNNs is complicated due to interpersonal variability and requires a long computational time. In this study, identical hyperparameters were applied to all the IO-CNNs for simplicity and efficiency. These hyperparameters were selected mainly by trial and error. Nevertheless, the CNN performance may be improved by individually tuning the CNN hyperparameters given the differences in aesthetic preferences between participants.

The use of CNNs in neuroscience to resemble biological brain computations is growing but remains controversial. As CNNs have many degrees of freedom, there are countless ways to project an input onto a desired output. Thus, CNNs can achieve high performance without mimicking the brain computations. Therefore, additional constraints derived from physiological observations should be included to reduce the wide diversity and help guiding the training of CNNs. In our study and various others [9, 5, 8, 44], a constraint has been related to the architecture mostly consisting of convolutions, inspired by single-cell receptive fields [45, 46]), and nonlinear activation, inspired by the rate of action potential firing [47]). A second constraint has been structural and provided by stacked convolutional layers that gradually increase the receptive field [48]. For example, stacking of two 3×3 convolutional layers yield an 5×5 receptive field. This approach resembles the increasing size of receptive fields in processing throughout the visual hierarchy [49]. In future work, we believe the further implementation of neuroscience-based techniques may help discover more appropriate CNN architectures for every brain region and hopefully provide insights about their integration to model whole-brain computations.

## Acknowledgements

This work was supported by Kao Corporation, JSPS KAKENHI 19K16252 and 20H05052 to TM; a grant from Brain/MINDS Beyond (AMED) to TM (grant number JP20dm0307031); a grant from JST-PRESTO to TM, JSPS KAKENHI 19K20390 to TQP, JSPS KAKENHI 18H05017 and 17H06033 to JC, a grant from Japan Agency for Medical Research and Development (AMED) to JC (grant number JP20dm0207086). Computational resources were provided by the Data Integration and Analysis Facility, National Institute for Basic Biology, and Research Center for Computational Science.

## Author contributions

Conceptualization: NS, JC; Methodology: TQP, JC, MT; Investigation: TQP, TY, HN, HKT,; Writing - Original Draft: TQP, JC; Writing - Review and Editing: TQP, TY, HTK, RU, TM, AKA, NS, JC; Supervision and Funding Acquisition: NS, JC.

## Methods

### Participants

Thirty-seven right-handed healthy participants [10 males; age 22.9±0.6 years; range, 18 – 32 years] with no history of neurological or psychiatric disorders were recruited from our locality via email for this study. They provided informed consent to participate in the study. The study was approved by the Ethical Committee of the National Institute for Physiological Sciences of Japan.

### Experimental procedure

The experiment consisted of two sessions, with a session of functional magnetic resonance imaging (fMRI) recordings on one day and a session of behavioral experiments on another day. The two experiments took place 29 to 147 days apart (79.3±7.0 days). The order of the experiments was balanced across participants.

### Visual stimuli

A total of 105,714 art images with price tags were downloaded from an art auction website (https://www.liveauctioneers.com/). From the images, 43,757 art pieces were actually traded. We excluded the art images with price tags above USD 5000 for most of the art images to be from non-widely known artists. This exclusion criteria aimed for the pricing of the participants to be based on their aesthetic preferences instead of the knowledge about an artist. We used 200 art images for the fMRI experiment, 400 images for the behavioral experiment, and 38,059 images for training the baseline convolutional neural network (CNN).

### fMRI experiment

In the fMRI session, 200 art images were randomly presented over five rounds per participant. The participants were asked to quote the prices that they were willing to pay for each art piece. In a trial, a photo of an oil painting was presented for 7 s followed by a blank screen for 1 s and by a price for 6 s. The participants could move the price on the window up or down by pressing the corresponding button. After a 4 s intertrial interval, the next photo was presented. The price data obtained from the fMRI experiment were used to optimize the CNN for each participant (IO-CNN).

### Behavioral experiment

The participants were asked to quote their target prices for a new set of 400 art pieces. Besides the pricing task, the participants completed a reliability check, in which they were asked to rank a separate set of 80 art pieces twice over a period of approximately 2 h. To measure reliability, we calculated the correlation between these two ratings per participant and then divided the participants into reliable and unreliable groups accordingly.

### Imaging parameters

The fMRI scans were collected using a 3.0 T fMRI system (Verio; Siemens Erlangen, Germany) with a 32-element phased-array head coil. T2*-weighted gradient echo-planar imaging (EPI) was used to obtain functional images using the following parameters: repetition time TR = 750 ms, echo time TE = 31 ms, flip angle of 55°, field of view of 192 mm, matrix size of 98×98, 72 slices with isotropic 2-mm voxels, and multiband echo-planar imaging with factor of 8. For anatomical imaging, T1-weighted three-dimensional magnetization-prepared rapid-acquisition with gradient echo sequence was employed using the following parameters: TR = 1800 ms, TE = 1.98 ms, flip angle of 9°,field of view of 256 mm; and voxel dimensions of 1.0×1.0×1.0*mm*.

### Data analysis

Standard image preprocessing was performed using the Statistical Parametric Mapping SPM12 package. The fMRI scans were realigned to correct for head motion. Each T1-weighted image per participant was co-registered with the mean image over all the functional images of that participant. Then, the functional images were segmented and normalized to the MNI 152 space using the unified segmentation–normalization tool in SPM12. To obtain precise voxels in the gray matter from each participant, the first echo-planar imaging volume in the first round was segmented into gray- and white-matter masks using the tissue probability map of SPM12. MELODIC (Multivariate Exploratory Linear Optimized Decomposition into Independent Components) was used to decompose functional data into spatially independent components, which were then manually labelled as either signal or noise [50]. FIX (FMRIB’s independent-component-analysis-based Xnoiseifier) was then applied for noise component regression. Each stimulus presentation was modeled as a separate event using the canonical function in SPM12. To visualize the imaging results, Connectome Workbench (https://www.humanconnectome.org/software/connectome-workbench) and MRIcron (https://people.cas.sc.edu/rorden/mricron/) were used.

### Representational Similarity Analysis

We used a searchlight analysis based on spheres of 4 mm in radius to include 33 voxels. A correlation matrix of art images was constructed per voxel and its 33 neighbors. Then, the matrix was described by multivariate linear regression of IO-CNN representational similarity matrices (Fig. 1).

To improve the estimation of regression coefficients, we employed ridge regression. The ridge parameter was determined based on a pilot study, and the same parameter was applied to every voxel for all the participants. Parameter β defines the coefficient (weight) for each IO-CNN representational similarity matrix. For group analysis, β maps for the respective layers were smoothed with a 4 mm full width at half maximum Gaussian kernel and then subject to one sample t-test across participants. In this group analysis, the statistical threshold for the spatial extent test on the clusters was set at *p*<0.05 with family-wise error corrected for multiple comparisons. The cluster-forming threshold was set at *p*<0.001 (uncorrected). To visualize the spatial configuration of brain–CNN correspondence, we assigned the layer with the highest t-value to each voxel, thus obtaining overall maps of brain–CNN correspondence (Figs. 2a, 2b, 4a, 4b, 5a and 6a). To analyze the brain–CNN correspondence in regions of interest (ROIs), we counted the number of significant voxels per CNN layer in each ROI (Fig. 3). Those numbers were divided by the number of significant voxels in the whole brain per CNN layer to obtain the proportion of significant voxels per layer.

### ROI definition

We used three atlases, namely, cortical area parcellation from resting-state correlations (Gordon atlas) (333 ROIs) [16], Human Connectome Project multimodal parcellation (360 ROIs), and anatomical automated labeling atlas (170 ROIs)[51]. The Gordon atlas was used to visualize spatial relations between the brain–CNN correspondence maps and the default mode network (Figs. 2a, 2b, 4a, 4b, 5a, and 6a). The multimodal parcellation was used to visualize the distribution of the layer-wise correspondence between the brain and CNN (Fig. 3) in a finer parcellation. The anatomical automated labeling atlas was used to define the occipital, temporal, parietal, and frontal cortices as well as the insula and cingulate gyri.

The occipital cortex consists of the calcarine fissure and surrounding cortex, cuneus, lingual gyrus, superior occipital lobe, middle occipital lobe, and inferior occipital lobe. The temporal cortex consists of the hippocampus, parahippocampus, amygdala, fusiform gyrus, Heschl gyrus, superior temporal gyrus, temporal pole, middle temporal gyrus, and inferior temporal gyrus. The parietal cortex consists of the postcentral gyrus, superior parietal gyrus, inferior parietal gyrus, supramarginal gyrus, angular gyrus, and precuneus. The frontal cortex consists of the precentral gyrus, superior frontal gyrus, middle frontal gyrus, inferior frontal gyrus, Rolandic operculum, supplementary motor area, olfactory cortex, superior frontal gyrus, gyrus rectus, and paracentral lobule.

### Comparison between principal gradient scores and brain–CNN correspondence maps

The principal gradient map was retrieved from the Neuroanatomy & Connectivity Lab website (https://www.neuroconnlab.org/data/index.html). To capture the correspondence between the hierarchical structure of the brain and CNN, the principal gradient scores per CNN layer were plotted (Figs. 2d). For statistical evaluation, we also calculated the correlations between the principal gradient scores per voxel at each CNN layer and the layer number (Fig. 2D, Tables **Error! Reference source not found**. and **Error! Reference source not found**.).

### CNN architecture

We used the VGG-16 architecture [48] with batch normalization and pretrained on ImageNet dataset [52]. The network was implemented using the PyTorch library (ver. 1.0.0) [53]. Originally, the network consists of 16 convolutional layers with batch normalization and two fully connected (FC) layers. Here, we replaced the last softmax function with three FC layers. Batch normalization and dropout regularization [54] were applied after each additional FC layer, except for the final layer which contained only one node for linear regression.. We applied Kaiming He initialization to the newly added hidden layer [55].

### Baseline CNN trained using transfer learning

A baseline CNN was created by training on a 38,059 art-price dataset (27,488, 3054, and 7517 samples for training, validation, and test sets respectively), retrieved from the art-auction website. During transfer learning, it is usual to freeze (i.e., exclude from training) all layers except for the last few layers, which process domain-general knowledge [56]. However, choosing the number of layers for freezing is not trivial and may lead to catastrophic forgetting if few layers are frozen or slow convergence if many layers are frozen. Therefore, we fine-tuned the models over three stages: training the three newly added FC layers, training all the FC layers, and finally training all the layers. Then, training terminated after the performance converged. This approach is similar to the gradual unfreezing method widely used in natural language processing[57].

### Individual CNN optimization

For each participant, transfer learning was applied to the CNN by using both the 200 art–price pairs for training and 400 art–prices pairs for validation and test. Adam optimization [58] was used with a weight decay of 0.00025.

All datasets were preprocessed as follows. First, we calculated the mean and standard deviation from the outputs of the baseline CNN on the 200 art–price pairs. Then, we normalized both the 200 and 400 art–price pairs to zero mean and unit standard deviation using the calculated mean and standard deviation. This method prevents the unwanted influence from different distributions between the baseline CNN output and art–price pairs. To prevent overfitting, the training data were augmented as follows. Each image was randomly translated in a range of 20% along the height and width directions. Then it was cropped for centering and resized to 224×224 pixels. This step was repeated five times on the fly. Therefore, the augmented training set (including the original dataset) was six times larger than the original training set. On the other hand, the 400 art–price pairs were equally distributed into the validation and test sets for performance testing. To keep the distribution of these sets identical, we divided the experimental data into 10 bins and randomly sampled the pairs for each bin.

For individual CNN optimization, we fine-tuned the models over four stages (Fig. 1b), and training terminated when the performance converged. For each of the first three training stages, we performed a nearly exhaustive search for hyperparameters, such as learning rate and batch size. To prevent exploding gradients, the gradient norm was truncated at 1.0. We tracked the mean squared error (MSE) on the validation set and recorded the parameters that minimized the MSE. Early stopping was used to terminate training after 10 epochs without performance improvement, and the maximum number of epochs for training was 200. The performance of each CNN was evaluated through the MSE between its output and the ground-truth values on the validation set.

In the fourth stage, we used Adam optimization with a high learning rate of 0.001 for restart. This strategy is inspired by the stochastic gradient descent with warm restarts (SGDR) [59], in which the learning rate is restarted after a fixed number of iterations. This strategy aims to leave any local optima and search for a better one (if any) while keeping track of the previous best performance. Again, early stopping was used, and the maximum number of epochs for training was 300. The parameters that minimized the MSE on the validation set were recorded.

The representational similarity matrices of all the IO-CNN layers were created from the outputs of the convolutional layers (13 layers) and FC layers (5 layers) on the 200 art–price pairs. We extracted and transformed the responses of each layer into an *n*_*images*_×*n*_*nodes*_ matrix. Then, we constructed the *n*_*images*_×*n*_*images*_ correlation matrix as the representational similarity matrix for that layer.

### Transfer learning of IO-CNNs based on ResNet, DenseNet, and Inception Network

We built IO-CNNs based on DenseNet, ResNet, and Inception Network using similar procedures to those mentioned above. As each architecture has several variants, we used ResNet-50, DenseNet-169, and Inception-v3 for convenience because they are already implemented in PyTorch as baseline CNNs pretrained on the ImageNet dataset [52]. The last softmax layer was replaced by three FC layers followed by batch normalization and dropout layers at the end of each layer except for the last one.

First, the three newly added FC layers were trained in each CNN, and then all the other layers were trained using the 38,059 art–price pairs. Then, for each subject, the baseline CNNs were trained using the individual valuation data over three stages: training the FC layers, training all the layers, and finally training all the layers with high learning rate for restart.

The total number of representational similarity matrices that were extracted differed between the architectures. For ResNet, we obtained 23 representations from the outputs of the bottleneck blocks (20 representations), and FC layers (3 representations). For Inception Network, 14 representations were obtained from the outputs of the inception blocks (11 representations) and FC layers (3 representations). For DenseNet, we obtained 10 representations from the dense blocks (4 representations), transition blocks (3 representations), and FC layers (3 representations).

