## Supplementary Materials for "Vision-to-value transformations in artificial neural networks and human brain"

Numer of figures: 3

Numer of tables: 3

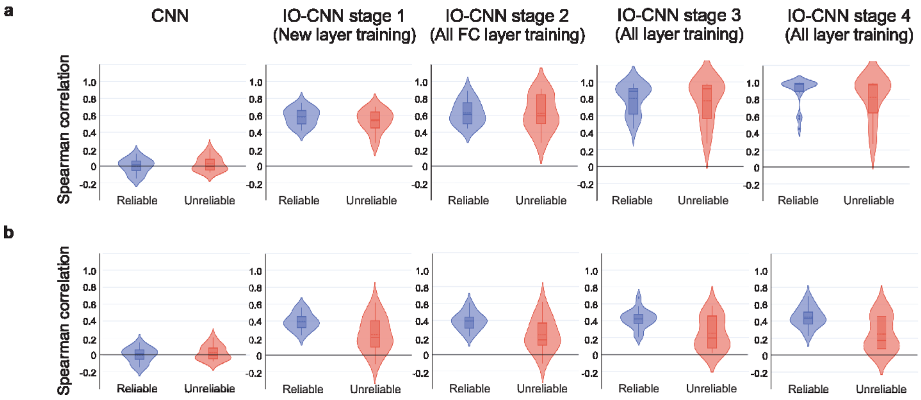


Figure S1: Comparison of prediction performance between the reliable and unreliable groups. (**a**) The change of prediction performance for the training dataset over individual optimization in the reliable (blue) and unreliable(red) groups. The violin plots represent the distributions of correlation. The boxes span the first to third quartiles. The solid line inside the boxes represents the median and the dashed line represents the mean. The dots represent outliners. (**b**) The change of prediction performance for the test dataset over individual optimization in the reliable (blue) and unreliable (red) groups.

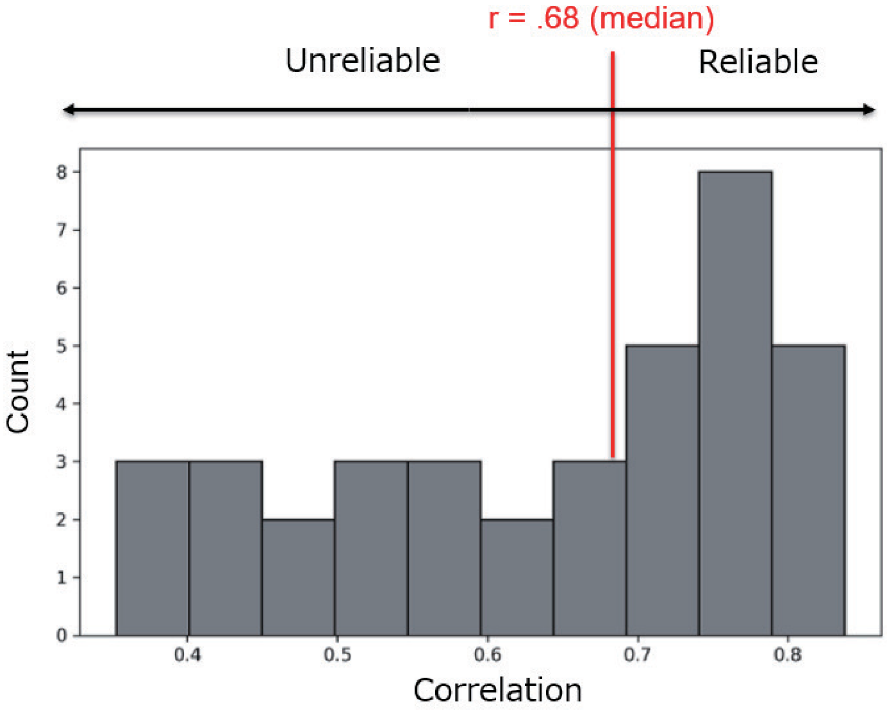


Figure S2: Distribution of correlation between participant's ratings for the same painting across an approximately 2-hour interval. The red line indices median (r=0.68).

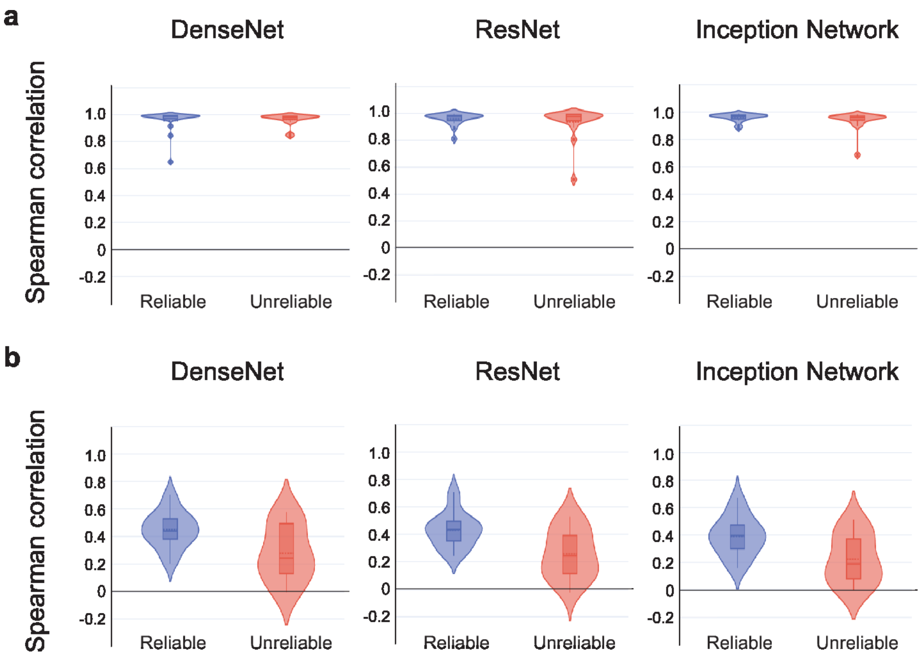


Figure S3: Prediction performance of different CNN architectures. (**a**–**b**) Correlation between model's predictions and ground truth values on train set (a) and test set (b), across 3 different CNN models (DenseNet, ResNet, Inception Network). The violin plots represent the distributions of correlation. The boxes span the first to third quartiles. The solid line inside the boxes represent the median, the dashed line represents the mean, and the dots represent outliers.

Table S1: List of ROIs abbreviations

| | |
|---|---|
| 44 | Area 44 |
| IFSp | Posterior Inferior Frontal Sulcus |
| p9-46v | Area posterior 9-46v |
| 46 | Area 46 |
| a9-46v | Area anterior 9-46v |
| IFSa | Anterior Inferior Frontal Sulcus |
| 45 | Area 45 |
| 47l | Area 47 lateral |
| d32 | Area dorsal 32 |
| 9m | Area 9 medial |
| 10d | Area 10 dorsal |
| p32 | Area posterior 32 |
| 10r | Area 10 rostral |
| a24 | Area anterior 24 |
| p24 | Area posterior 24 |
| 23c | Area 23c |
| 31a | Area 31a |
| PCV | PreCuneus Visual Area |
| 31pd | Area 31pd |
| 7m | Area 7 medial |
| v23ab | Area ventral 23 a+b |
| 31pv | Area 31p ventral |
| d23ab | Area dorsal 23 a+b |
| 23d | Area 23d |

Table S2: Correlation between PG scores and corresponding layer number at each stage of the individual optimization of CNN (VGG-based model). *: p <.001 (FWE)

| | CNN | IO-CNN Stage 1 | IO-CNN Stage 2 | IO-CNN Stage 3 | IO-CNN Stage 4 |
|---|---|---|---|---|---|
| Reliable | *0.06 | *0.14 | *0.15 | *0.33 | *0.36 |
| Unreliable | *0.03 | *0.07 | *0.04 | *0.21 | *0.24 |

Table S3: Correlation between PG scores and CNN layers for three popular CNN architectures. *: p <0.001 (FWE)

| | ResNet | Inception Network | DenseNet |
|---|---|---|---|
| Reliable | *0.30 | *0.41 | *0.38 |
| Unreliable | *0.23 | *0.23 | *0.26 |
